# ZBP1-mediated macrophage necroptosis inhibits ASFV replication

**DOI:** 10.1101/2023.04.25.538262

**Authors:** Keshan Zhang, Yu Hao, Bo Yang, Jinke Yang, Dajun Zhang, Xing Yang, Xijuan Shi, Dengshuai Zhao, Lingling Chen, Wenqian Yan, Yi Ru, Zixiang Zhu, Xiaodong Qin, Huanan Liu, Fan Yang, Dan Li, Hong Tian, Tao Feng, Jianhong Guo, Jijun He, Xiangtao Liu, Haixue Zheng

## Abstract

African swine fever (ASF) is an infectious disease characterized by hemorrhagic fever, highly pathogenic, and severe mortality in domestic pigs caused by the African swine fever virus (ASFV). ASFV is a large DNA virus and primarily infects porcine monocyte macrophages. The interaction between ASFV and host macrophages is the major reason for gross pathological lesions caused by ASFV. Necroptosis is an inflammatory programmed cell death and plays an important immune role during virus infection. However, whether and how ASFV induces macrophage necroptosis and what effect of necroptosis signaling on host immunity and ASFV infection remains unknown. This study uncovered that ASFV activates the necroptosis signaling in the spleen, lung, liver, and kidney from ASFV-infected pigs. And macrophage necroptosis also was induced by ASFV infection in vitro. Further evidence showed that the macrophage necroptosis is independent of TNF-α-RIPK1 or TLR-TRIF pathway but depends on Z-DNA binding protein 1 (ZBP1). ASFV infection upregulates the expression of ZBP1 and RIPK3 to consist of the ZBP1-RIPK3-MLKL necrosome and further activates macrophage necroptosis. Subsequently, multiple Z-DNA sequences were predicted to be present in the ASFV genome. And the Z-DNA signals were further confirmed to be present and colocalized with ZBP1 in the cytoplasm and nucleus of ASFV-infected cells. Moreover, ZBP1-mediated macrophage necroptosis caused the extracellular release of proinflammatory cytokines TNF-α and IL-1β induced by ASFV infection. Finally, we demonstrated that ZBP1-mediated necroptosis signaling significantly inhibits ASFV replication in host macrophages. Our findings uncovered a novel mechanism by which ASFV induces macrophage necroptosis by facilitating Z-DNA accumulation and ZBP1 necrosome assembly, providing significant insights into the pathogenesis of ASFV infection.

**Importance:** ASFV infection causes an acute, febrile, hemorrhagic, and severely lethal disease in swine, which seriously threatens the global porcine industry. Understanding the interaction mechanism between ASFV and host macrophages during infection is essential for elucidating the pathogenesis of ASFV. To our knowledge, this study revealed the interaction mechanism between ASFV infection and host macrophage necroptosis. The results showed that ASFV infection induces macrophage necroptosis through ZBP1 activation instead of the TNF-α-RIPK1 or TLR-TRIF pathway. ASFV infection promotes Z-DNA accumulation, which causes ZBP1-RIPK3-MLKL necrosome assembly and macrophage necroptosis. The ZBP1-mediated necroptosis signaling facilitates the extracellular release of proinflammatory cytokines, inhibiting ASFV replication in host macrophages. This study found a new interaction mechanism between ASFV and host macrophages, which may help understand the pathogenesis caused by ASFV infection.

## Introduction

African swine fever (ASF), caused by the African swine fever virus (ASFV), is an extremely contagious and highly fatal disease in domestic pigs and wild boars. The spread of ASF worldwide has severely impacted the global pig industry over the past century (1). In 2018, ASF broke out in China and spread rapidly throughout the country, causing great economic losses and posing a serious threat to Chinese pork safety and supply (2, 3). Unfortunately, the absence of effective vaccines or drugs for ASFV remains, which caused ASF prevention only relying on the rapid culling of susceptible animals and strict epidemic control measures (4).

ASFV, the only member of the *Asfarviridae* family, is a large, enveloped, double-stranded (ds) DNA virus (5). The dsDNA genome length of ASFV ranges from 170 to 194 kb encoding more than 150 proteins indispensable to viral replication, transcription, and host immune evasion (6). ASFV mainly infects porcine monocyte macrophages, and the interaction between ASFV and porcine macrophages is the main cause of ASFV pathogenesis (7-9). However, the interaction between ASFV and porcine macrophages has not been distinctly clarified, even with much effort.

Programmed cell death (PCD) is an effective immune mechanism by which host cells restrict pathogen infection. ASFV infection causes the macrophages’ apoptosis dependent on caspases 3, 7, 9, and 12 (10, 11). And tumor necrosis factor α (TNF-α)-dependent extrinsic apoptosis was induced in porcine alveolar macrophages (PAMs) infected with ASFV (8). ASFV has also evolved multiple viral proteins, maintaining its fast replication by interfering with host apoptosis machinery. For example, A179L, a Bcl-homologue encoded ASFV, can inhibit mitochondrial apoptosis by interacting with the BH3 domain of pro-apoptotic Bcl-2 family proteins (12). And ASFV A224L, an IAP-like protein, suppresses the proteolytic cleavage of caspase 3 by antagonizing TNF-α signaling (13). In addition to apoptosis, pyroptosis, an inflammatory PCD characterized by the activation of the inflammasome, maturation of inflammatory cytokines, and plasma membrane damage, is also hijacked by ASFV (14). ASFV-encoded pS273R relieved the pyroptotic cell death by inactivating execution protein gasdermin D (GSDMD) (15). And the interaction between ASFV pH240R and host NLRP3 inhibits macrophage pyroptosis by limiting the NLRP3 oligomerization and IL-1β production (16).

Necroptosis, a caspase-independent inflammatory PCD similar to pyroptosis, induces intense inflammation by promoting the damage of the plasma membrane and releasing the inflammatory cytokines and damage-associated molecular patterns (DAMPs) into the cytoplasm (17). Initiation of necroptosis signaling requires the recruitment of receptor-interacting protein (RIP) homotypic interaction motif (RHIM)-containing RIP kinase 3 (RIPK3) to one of the other three RHIM-containing proteins, including receptor-interacting protein kinase 1 (RIPK1), TIR-domain-containing adapter-inducing interferon (TRIF) and Z-conformation nucleic acid (Z-NA) binding protein 1 (ZBP1) (18). After recruiting to other RHIM-containing proteins, RIPK3 phosphorylates and interacts with mixed lineage kinase domain-like protein (MLKL) to consist necrosome. Phosphorylated MLKL in necrosome polymerizes on the cytoplasmic membrane, forming membrane pores and driving plasma membrane leakage (19, 20). TNF-α induced RIPK1-RIPK3-MLKL necrosome, Toll-like receptor (TLR) 3/4 mediated TRIF-RIPK3-MLKL necrosome, and ZBP1-RIPK3-MLKL necrosome play a significant role in infectious diseases, such as herpesvirus (HSV), vaccinia virus (VACV), influenza A virus (IAV), and enteropathogenic *Escherichia coli* infection (21).

Porcine TNF-α is abundantly secreted in porcine serum and cell culture supernatants after ASFV infection (7, 8, 22). And ASFV-encoded A179L has also enhanced RIPK1-dependent necroptosis induced by TNF-α in L929 cells (23). Moreover, TLR3/4 is activated to promote IL-1β expression in ASFV-infected PAMs (24). These past discoveries suggest ASVF-induced necroptosis signaling is possible via TNF-α-RIPK1 or TLR3/4-TRIF pathways. However, it remains unknown whether and how ASFV triggers the necroptosis signaling and what effect of ASFV-induced necroptosis on host immunity and ASFV infection. Thus, further research on the interaction between ASFV and macrophage necroptosis is necessary.

This study found that ASFV efficiently activates the necroptosis signaling in the porcine spleen, lung, liver, and kidney. We also confirmed that ASFV induces porcine macrophage necroptosis in *vitro*. Next, we demonstrated that ASFV-induced macrophage necroptosis was independent of TNF-α-RIPK1 or TLR-TRIF pathways but depended on upregulated ZBP1 in porcine macrophages. Furthermore, we uncovered that ASFV infection induces the formation of Z-conformation DNA (Z-DNA) to activate ZBP1-mediated necroptosis, causing the extracellular release of proinflammatory cytokines in porcine macrophages. ZBP1-mediated necroptosis signaling restricts ASFV replication in porcine macrophages. Our findings uncover an interaction mechanism between ASFV and porcine macrophage necroptosis that further elucidates the pathogenesis of ASFV.

## Results

### ASFV infection activated the necroptosis signaling in *vivo*

To explore whether ASFV activates necroptosis signaling, tissue samples including spleen, lung, liver, and kidney were taken from health or ASFV-infected pigs. Then, the expression and phosphorylation levels of RIPK3 and MLKL in these tissue samples, the hallmark of necroptosis, were examined using a Western blot. The results showed that the expression and phosphorylation levels of RIPK3 and MLKL were undetectable or faint in healthy pigs’ spleen, lung, liver, and kidney. However, the expression and phosphorylation levels of RIPK3 and MLKL were significantly upregulated in ASFV-infected spleen, lung, liver, and kidney (**Fig. 1A-D**). Therefore, necroptosis signaling was efficiently activated by ASFV in pigs’ spleen, lung, liver, and kidney.

**FIG 1.**
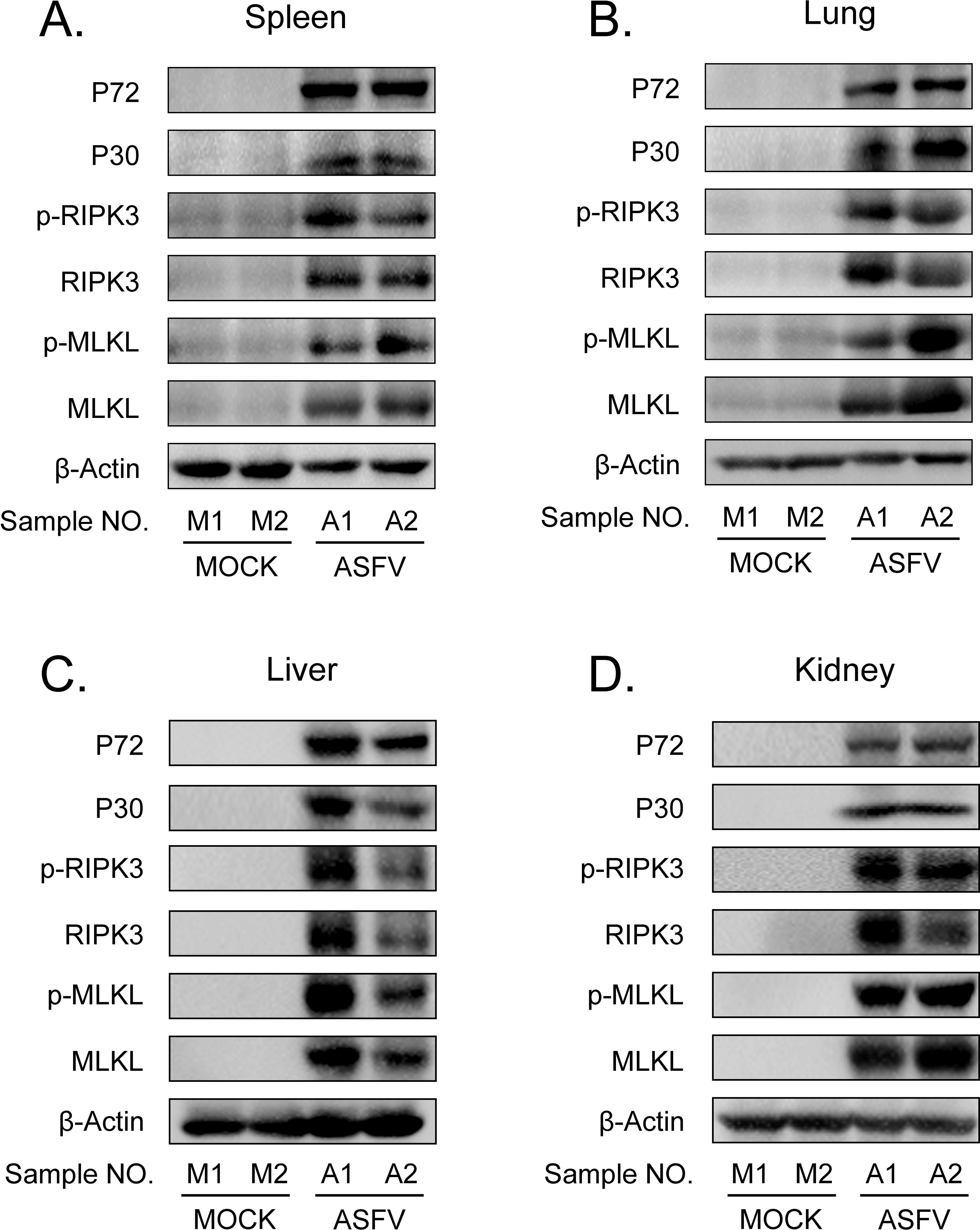
ASFV infection activates the necroptosis signaling in *vivo*. (A-D) Western blot detects the protein expression and protein phosphorylation level of RIPK3 and MLKL in the spleen (A), lung (B), liver (C), and kidney (D) collected from two healthy pigs (M1 and M2) and two ASFV-infected pigs (A1 and A2). ASFV structure proteins P72 and P30 were detected to confirm ASFV infection. And β-actin was selected as the internal reference protein.

### ASFV infection efficiently induces macrophage necroptosis in *vitro*

Next, the necroptosis signaling in ASFV-infected primary PAMs and porcine bone marrow-derived macrophages (BMDMs) was analyzed. ASFV replicates efficiently in PAMs and BMDMs, and its proliferation kinetics are shown in Figure S1.

After infecting PAMs and BMDMs with ASFV in dose-dependent or time-dependent ways, the expression and phosphorylation levels of RIPK3 and MLKL were analyzed using Western blot cytoplasmic membrane integrity of ASFV-infected macrophages also was determinated by cytotoxicity analysis.

The results indicated that the expression of RIPK3 and the phosphorylation levels of RIPK3 and MLKL were significantly elevated in ASFV-infected PAMs and BMDMs in a dose-and time-dependent manner (**Fig. 2A, 2C, 2E, and 2G**). That is similar to the data from ASFV-infected porcine tissues. Differently, MLKL is expressed constructively in PAMs and BMDMs. Besides, the release of cytoplasmic lactate dehydrogenase (LDH) in the supernatants of ASFV-infected PAMs and BMDMs significantly increased in a dose-and time-dependent manner, indicating that the cytoplasmic membrane of host macrophages was damaged during ASFV infection (**Fig. 2B, 2D, 2F, and 2H**). Together, these results confirmed that ASFV infection induces macrophage necroptosis in *vitro*.

**FIG 2.**
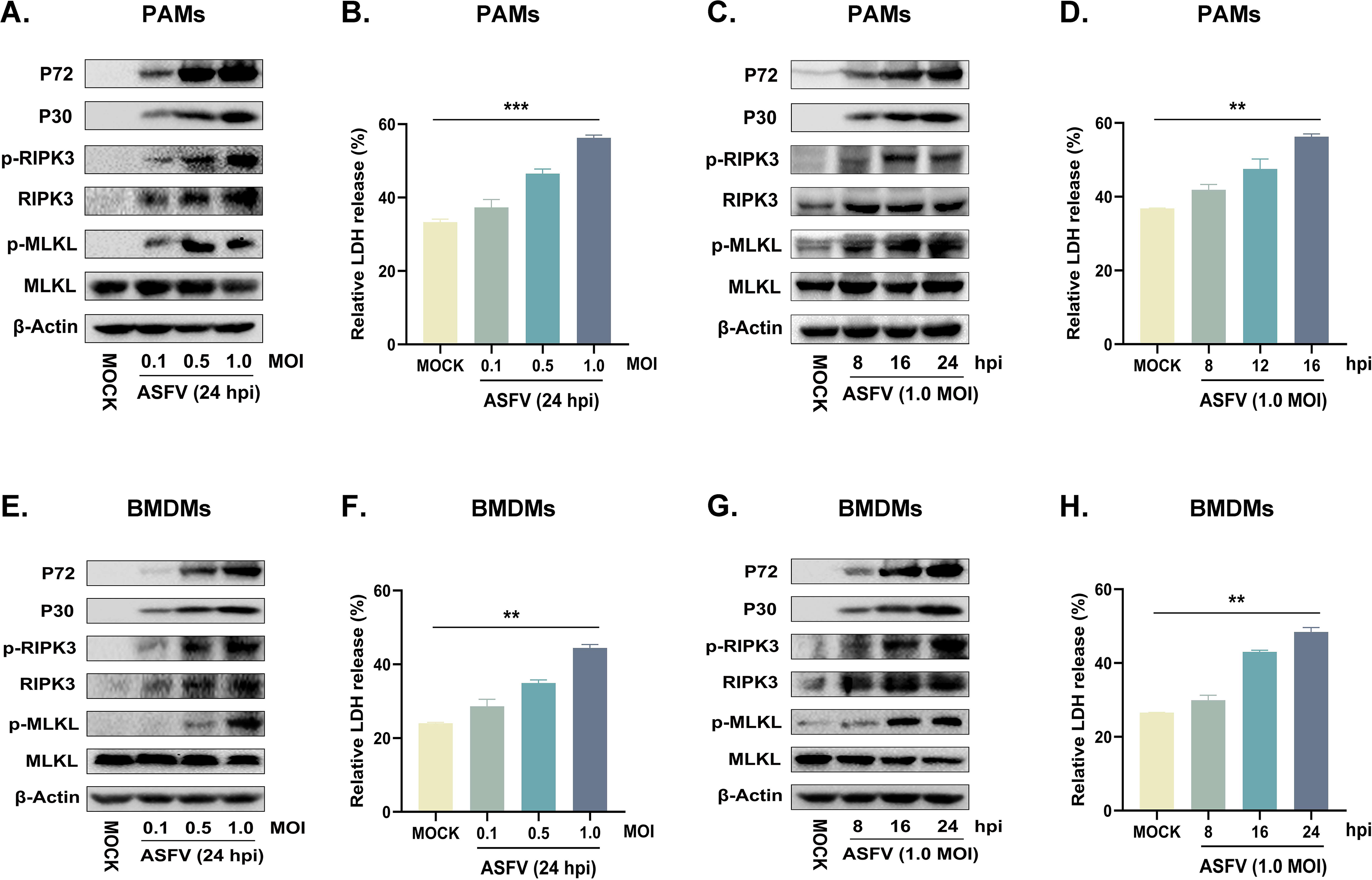
ASFV infection activates the necroptosis signaling in PAMs and BMDMs in *vitro*. PAMs (A and B) or BMDMs (E and F) were infected with ASFV at an MOI of 0.1, 0.5, or 1 for 24 h, or PAMs (C and D) or BMDMs (G and F) were infected with ASFV at an MOI of 1.0 for 8, 16 or 24 h. (A, C, E, and G) The protein expression and protein phosphorylation level of RIPK3 and MLKL in cell lysates (Lys) were examined by Western blot with the indicated antibodies. (B, D, F, and H) The relative LDH release in the supernatants was determined by cytotoxicity assay, and the level of LDH in fully lysed cells was defined as one hundred percent. Data shown are means ± SD. Statistical significance was analyzed by Student’s t-test. \**P* < 0.05; \*\**P* < 0.01; \*\*\**P* < 0.001; \*\*\*\**P* < 0.0001.

### ASFV-induced necroptosis signaling is independent of the TNF-α-RIPK1 pathway in macrophages

ASFV infection induces the high-level secretion of porcine TNF-α (7, 8, 22). And RIPK1-dependent necroptosis can be enhanced by overexpressing ASFV A179L (23). We sought to determine whether ASFV-induced macrophage necroptosis depends on the TNF-α-RIPK1 pathway. PAMs and BMDMs were treated with recombinant interferon-β (IFN-β) to induce the expression of RIPK3, then cotreated with TSZ (T: TNFα; S: SM-164; Z: Z-VAD-FMK) as the positive necroptosis control dependent on TNF-α-RIPK1 pathway. Necrostatin-1 (Nec-1, an inhibitor of kinase activity of RIPK1) was added to inhibit TNF-α-RIPK1 signaling in macrophages. And inhibitors GSK′872 (an inhibitor of the kinase activity of RIPK3) or GW806742X (an inhibitor of MLKL) were used to suppress necroptosis signaling activated via any pathway.

As shown in **Fig 3A and 3B**, the treatment of GSK′872 or GW806742X resulted in the obvious inhibition of MLKL phosphorylation and the release of LDH in ASFV-infected or TSZ-treated PAMs. While Nec-1 treatment significantly inhibited the phosphorylation of RIPK3 and MLKL and the release of LDH in TSZ-treated PAMs, a similar effect of Nec-1 was not observed in ASFV-infected PAMs. Similar to data from ASFV-infected PAMs, Nec-1 also failed to alleviate the phosphorylation of RIPK3 and MLKL and the release of LDH in ASFV-infected BMDMs (**Fig. 3C and 3D**). Thus, the TNFα-RIPK1 pathway is dispensable for ASFV-induced macrophage necroptosis.

**FIG 3.**
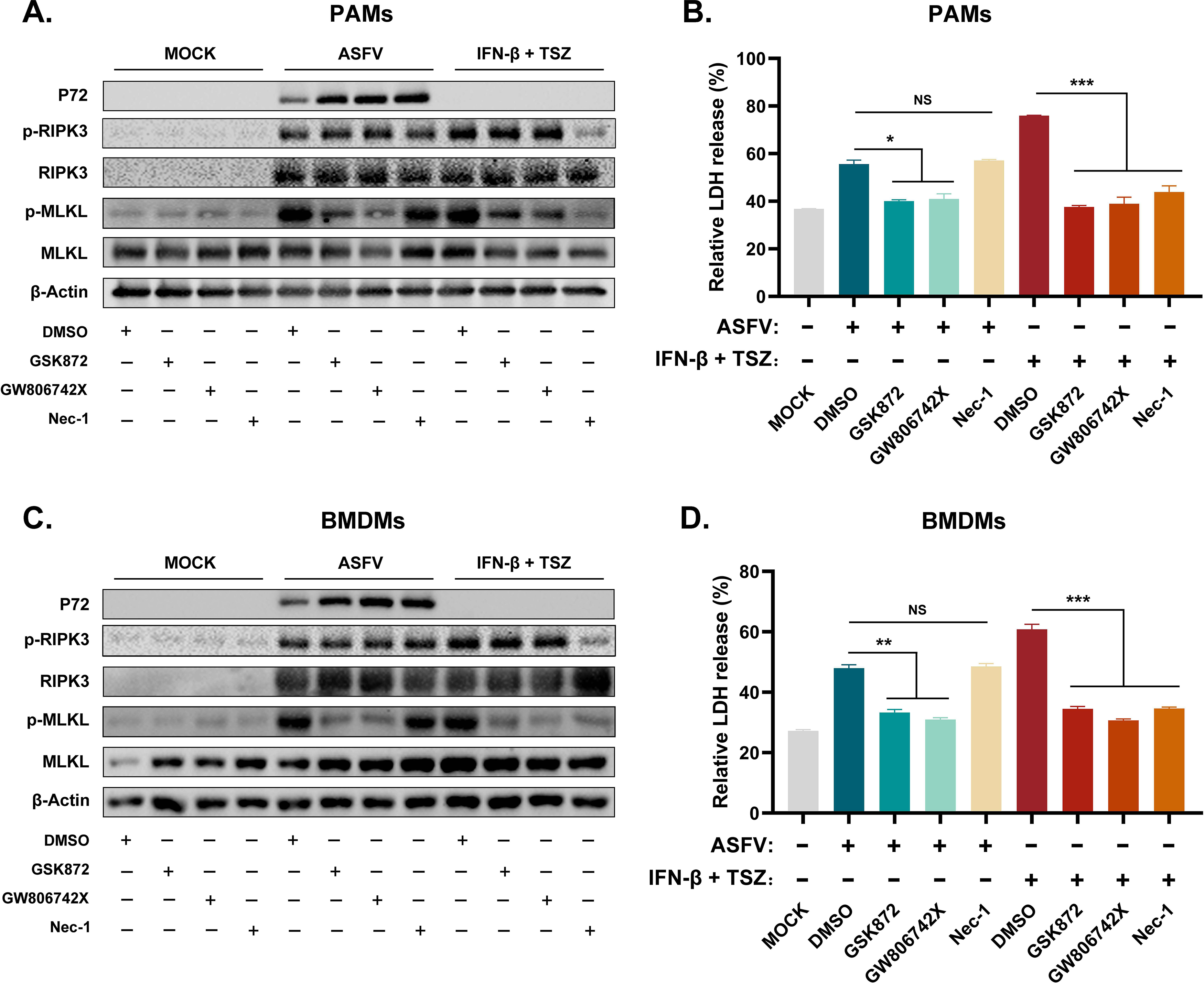
The ASFV-induced necroptosis signaling depends on the TNF-α-RIPK1 pathway in PAMs and BMDMs. PAMs (A and B) or BMDMs (C and D) were infected with ASFV at 1.0 MOI for 24 h to induce necroptosis, while cells without treatment (MOCK) were used as the negative control. PAMs (A and B) and BMDMs (C and D) were pretreated with IFN-β (1000 U/ml) for 8 h to induce the expression of RIPK3 and then were cotreated with TSZ (T: 100 ng/ml recombine porcine TNF-α; S: 50 nM SM-164; Z: 50 μM Z-VAD-FMK) for 16 h to activate the TNF-α-RIPK1 pathway-dependent necroptosis as the positive control. During infecting with ASFV or treatment with TSZ, cells were treated with RIPK3 inhibitor GSK’872 (1 μM) or MLKL inhibitor GW806742X (1 nM) to inhibit the necroptosis signaling. RIPK1 inhibitor Nec-1 (50 μM) was used to inhibit the necroptosis signaling dependent on the TNF-α-RIPK1 pathway. (A and C) The protein expression and phosphorylation level of RIPK3 and MLKL in cell lysates (Lys) were examined by Western blot. (B and D) The relative LDH release in the supernatants was determined by cytotoxicity assay. Data shown are means ± SD. Statistical significance was analyzed by Student’s t-test. \**P* < 0.05; \*\**P* < 0.01; \*\*\**P* < 0.001; \*\*\*\**P* < 0.0001.

### ASFV-induced necroptosis signaling is independent of the TLR-TRIF pathway in macrophages

TLR3 (an innate immune receptor sensing dsRNA) and TLR4 (an innate immune receptor sensing LPS) are important to the expression of IL-1β in ASFV-infected PAMs (24). We further investigated whether ASFV-induced macrophage necroptosis is dependent on the TLR3/4-TRIF pathway. PAMs and BMDMs were cotreated with IFN-β, LPS plus Z-VAD-FMK as the positive necroptosis control dependent on the TLR4-TRIF pathway. Pepinh-TRIF (TFA, a peptide blocking TLR-TRIF interaction) was added to inhibit TLR-TRIF signaling in macrophages.

The results showed that TFA treatment significantly inhibited the phosphorylation of RIPK3 and MLKL and the release of LDH in LPS-induced PAMs and BMDMs necroptosis. However, TFA still did not suppress the necroptosis signaling in PAMs and BMDMs induced by ASFV infection (**Fig. 4**). Therefore, these results indicated that ASFV-induced macrophage necroptosis is independent of the TLR-TRIF pathway.

**FIG 4.**
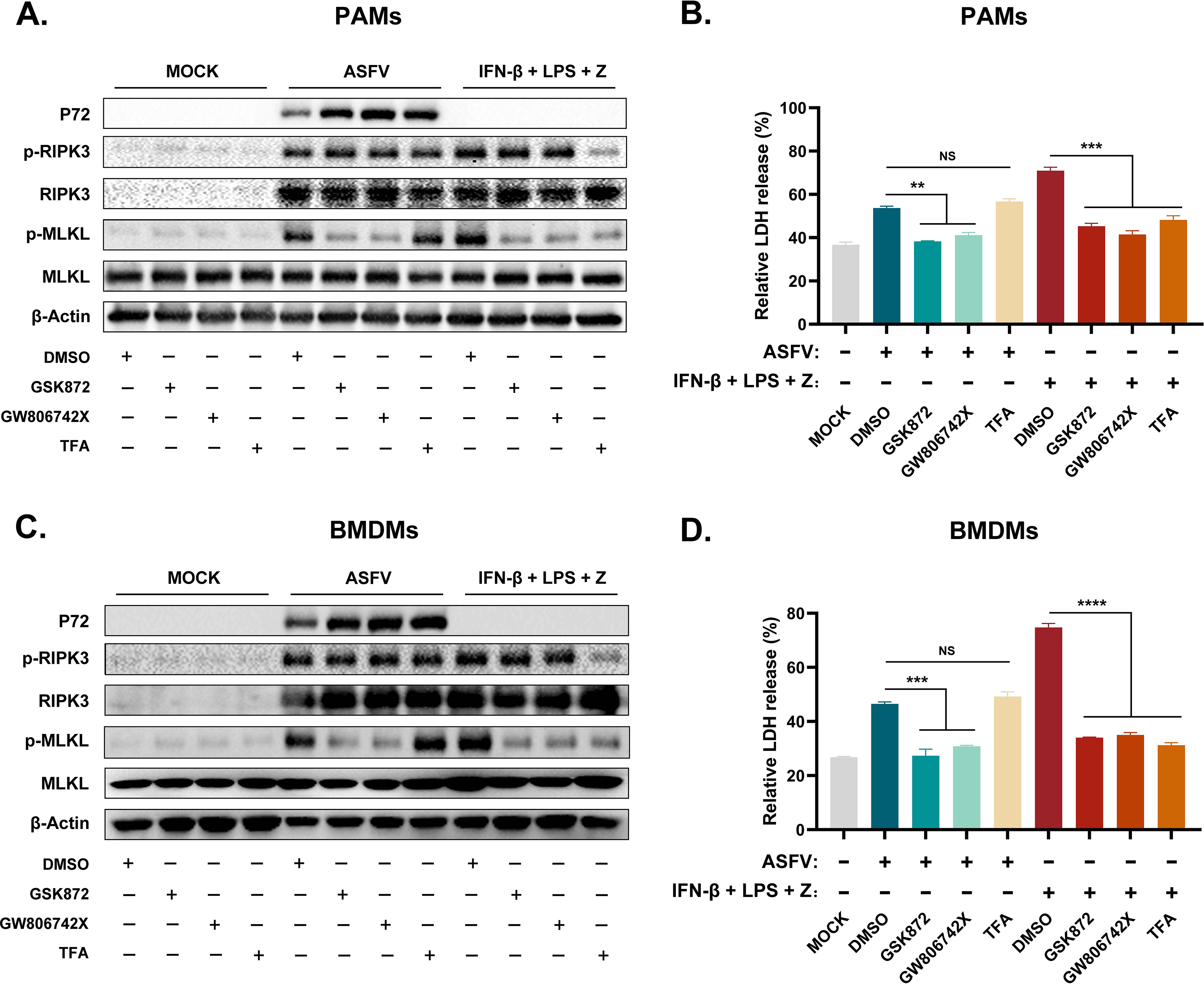
The ASFV-induced necroptosis signaling depends on the TLR-TRIF pathway in PAMs and BMDMs. PAMs (A and B) or BMDMs (C and D) were infected with ASFV at 1.0 MOI for 24 h to induce necroptosis, while cells without treatment (MOCK) were used as the negative control. PAMs (A and B) and BMDMs (C and D) were pretreated with IFN-β (1000 U/ml) for 8 h to induce the expression of RIPK3 and then were treated with LPS (100 ng/ml) plus Z-VAD-FMK (50 μM) for 24 h to activate the TLR4-TRIF pathway-dependent necroptosis as the positive control. During infecting with ASFV or treatment with LPS plus Z-VAD-FMK, cells were treated with GSK’872 (1 μM) or GW806742X (1 nM) to inhibit the necroptosis signaling. TFA (the interaction inhibitor between TLR and TRIF) (40 μM) inhibited the necroptosis signaling dependent on the TLR-TRIF pathway. (A and C) The protein expression and phosphorylation level of RIPK3 and MLKL in cell lysates (Lys) were examined by Western blot. (B and D) The relative LDH release in the supernatants was determined by cytotoxicity assay. Data shown are means ± SD. Statistical significance was analyzed by Student’s t-test. \**P* < 0.05; \*\**P* < 0.01; \*\*\**P* < 0.001; \*\*\*\**P* < 0.0001.

### ZBP1 necrosome is essential for ASFV-induced macrophage necroptosis

In addition to RIPK1 and TRIF, ZBP1 (an IFN-stimulated gene) is the third RHIM-containing host protein that recruits RIPK3 to activate necroptosis signaling. Thus, the effect of ZBP1 on ASFV-induced macrophage necroptosis was further explored. Our past research on transcriptomics, and proteomics of ASFV-infected host macrophages revealed that ZBP1 upregulates with interferon expression in macrophages during ASFV infection (**Fig. 5A**) (25, 26). The upregulation of ZBP1 was further confirmed by quantitative reverse transcriptase PCR (qRT-PCR) and Western blot analysis on ASFV-infected PAMs (**Fig. 5B and 5C**). Besides, compared to uninfected pigs, the expression of ZBP1 was significantly increased in the spleen, lung, liver, and kidney derived from ASFV-infected pigs (**Fig. S2**). To further explore the connection between ZBP1 and macrophage necroptosis during ASFV infection, endogenous co-immunoprecipitation (Co-IP) assays were conducted with ASFV-infected PAMs using an anti-ZBP1 antibody. PAMs without any treatment (MOCK) were used as a negative control, and IFN-β-stimulated PAMs were used as infection-free controls expressing ZBP1 and RIPK3.

**FIG 5.**
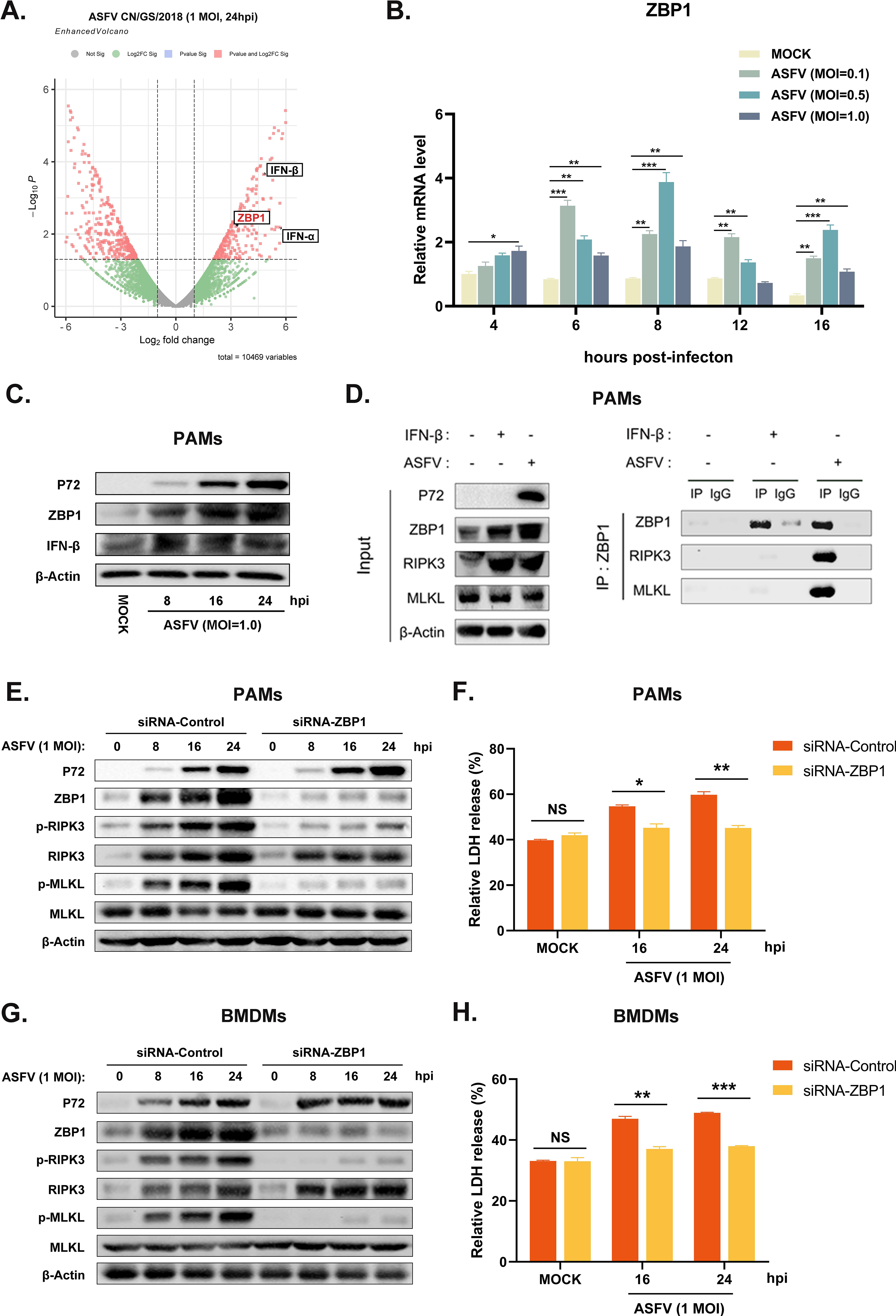
ASFV promotes the assembly of ZBP1 necrosome to induce macrophage necroptosis. (A) Transcriptomic analysis of ASFV-infected (MOI = 1.0) PAMs harvested at 24 hpi. Volcanic map profiling the mRNA expression level of ZBP1 and IFN-Ⅰ upon ASFV infection. Red dots indicated the differentially expressed genes (Log2 fold change>2 or <-2), and the left and right sides of the graph indicated the downregulated and upregulated genes, respectively. The transcription (B) and expression (C) levels of ZBP1 in PAMs infected with ASFV were detected by qRT-PCR and Western blot. GAPDH was selected as the internal reference gene of qRT-PCR. (C) PAMs were infected with ASFV at 1.0 MOI for 24 h, or treated with IFN-β (1000 U/ml) for 16 h, subjected to Co-IP using an anti-ZBP1 antibody. Nonspecific mouse IgG was used as a negative control. And the presence of RIPK3 and MLKL proteins was analyzed through immunoblotting using anti-RIPK3 and anti-MLKL antibodies. PAMs (E and F) or BMDMs (G and H) were infected with ASFV at 1.0 MOI for 0, 8, 16, or 24 h. Before infecting with ASFV, PAMs and BMDMs were transfected with siRNA targeting porcine ZBP1 to knock down ZBP1 expression or with nonspecific siRNA as a negative control. The knockdown efficiency was confirmed by Western blot using an anti-ZBP1 antibody. (E and G) The expression and phosphorylation level of RIPK3 and MLKL in cell lysates (Lys) were examined by Western blot. (F and H) The relative LDH release in the supernatants was determined by cytotoxicity assay. Data shown are means ± SD. Statistical significance was analyzed by Student’s t-test. \**P* < 0.05; \*\**P* < 0.01; \*\*\**P* < 0.001; \*\*\*\**P* < 0.0001.

The results of Western blot showed that ZBP1 and RIPK3 both highly expressed in IFN-β-stimulated and ASFV-infected PAMs. Interestingly, while ZBP1 not interacted with RIPK3 and MLKL in MOCK or IFN-β-stimulated PAMs, there is significant interaction between ZBP1, RIPK3, and MLKL was observed in PAMs upon with ASFV infection (**Fig. 5D**). These results demonstrated that ZBP1 necrosome composed of ZBP1, RIPK3, and MLKL was successfully assembled during ASFV infection.

To further confirmed the importance of ZBP1 on ASFV-induced macrophage necroptosis, the upregulated expression of ZBP1 in PAMs and BMDMs was knocked down by transfecting siRNA targeting ZBP1 mRNA. Then, the ASFV-induced necroptosis signaling was analyzed. Western blot results indicated that the phosphorylation of RIPK3 and MLKL was inhibited when ZBP1 upregulation disappeared in PAMs and BMDMs infected with ASFV (**Fig. 4E and 4G**). Moreover, ASFV-induced release of LDH also was dramatically alleviated in ZBP1-knocking down PAMs and BMDMs (**Fig. 4F and 4H**). These findings demonstrated that ASFV infection upregulates the ZBP1 expression to consist of necrosome and induce macrophage necroptosis.

### Z-DNA form and are recognized by ZBP1 in ASFV-infected PAMs

Sensing the virus-derived Z-nucleic acid (Z-NA) is critical for ZBP1 to activate necroptosis during virus infection (27). Considering that the ZBP1 necrosome is only assembled with ASFV infection (**Fig. 5D**), we hypothesized that ASFV infection generates the viral-derived Z-NA to activate ZBP1. Thus, the propensity to form Z-DNA sequences in the ASFV genome was first analyzed to test this hypothesis. Continuous (GC)_n_ in double-stranded nucleic acids is likely to form the Z-conformations, and the Z-forming potential of ASFV can be scored computationally by the Z-HUNT3 algorithm (28). The results of the Z-HUNT3 algorithm showed that the proportion of G and C in the ASFV CN/GS/2018 genome is about 40%, but the ASFV genome was predicted to contain multiple potential Z-DNA sequences (**Fig. 6A and 6B**). Past studies have confirmed that anti-Z-NA antibody (Z22) can recognize virus-derived Z-NA as the activated ligand recognized by the ZBP1 sensor. The same antibody was used to verify whether ASFV infection generates Z-NA generation. As shown in **Fig 6C**, the specific Z-NA signals were clearly observed in ASFV-infected PAMs with labeled anti-P30 and anti-Z-NA antibody. Because Z-NA antibody only recognize the Z conformation of double-stranded NA but not tell whether it is Z-DNA or Z-RNA, ASFV-infected PAMs were further treated with RNase A or DNase I. IFA results showed that the treatment with RNase A no significant effect on the accumulation of Z-NA signal (**Fig. 6D and 6F**), while DNase I treatment strongly reduced the Z-NA signals (**Fig. 6E and 6F**), suggesting that the detected Z-NA within ASFV-infected PAMs was Z-DNA. Besides, we found that the Z-DNA signals were distributed in both the cytoplasm and nucleus of PAMs infected with ASFV (**Fig. 6C and 6D**).

**FIG 6.**
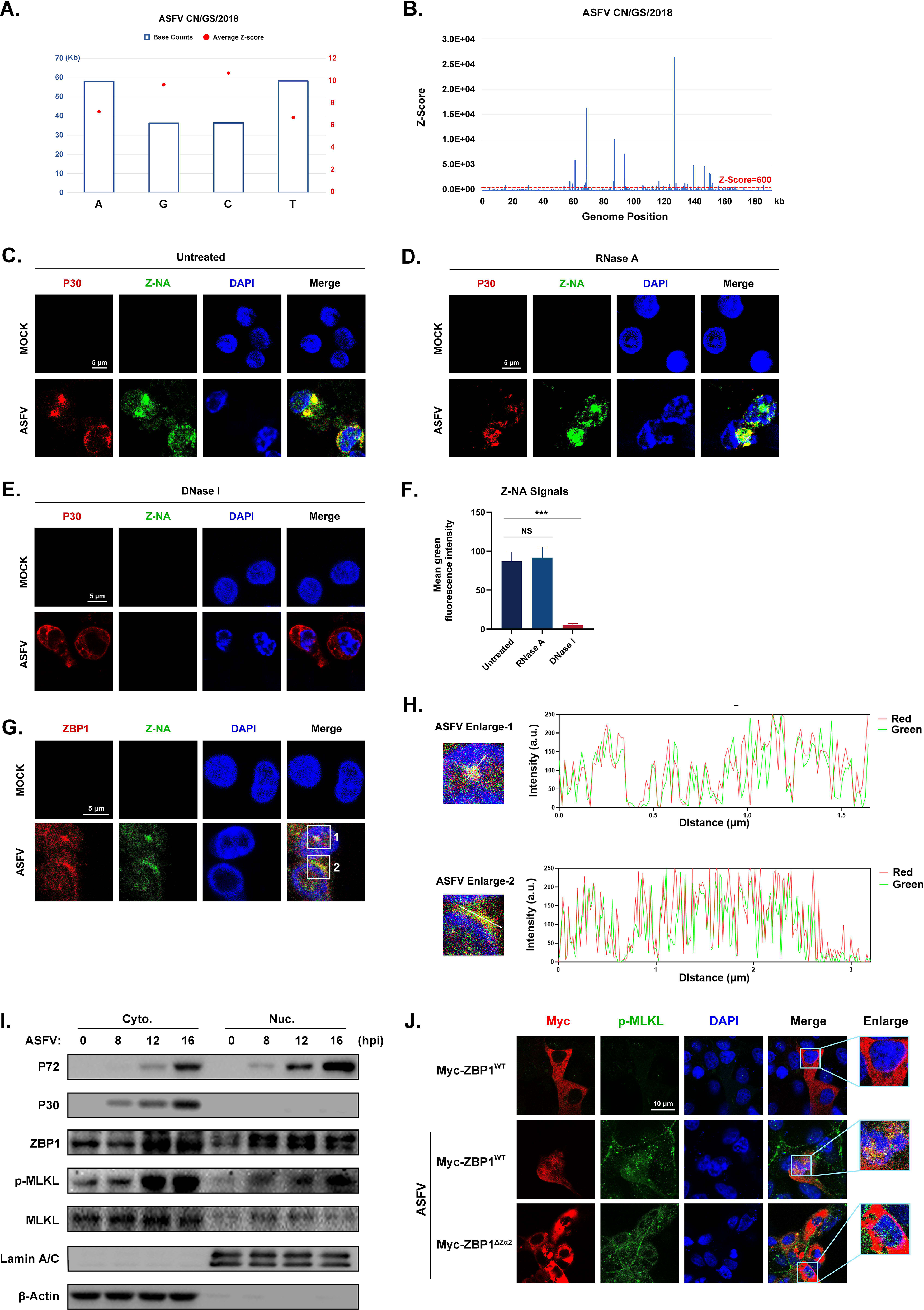
Accumulation of Z-DNA in ASFV-infected cells. (A) The counts and average Z-score of adenine (A), thymine (T), guanine (G), and cytidine (C) in the ASFV CN/GS/2018 genome. (B) The potential Z-DNA sequence in the ASFV genome was analyzed using the Z-HUNT 3 program, which scores the propensity of sequences to flip to the left-handed helical Z-conformation. The red dotted line indicated a Z-score of 600. A Z-score greater than 600 indicates that the sequence tends to form the Z-conformation under physiological conditions. (C-E) ASFV CN/GS/2018 (MOI = 1.0) infected or uninfected (mock) PAMs were fixed and permeabilized at 16 hpi and then either untreated (C) or treated with RNase A (1 mg/ml) (D) or DNase I (5 U/ml) (E). Cells were co-stained with antibodies against Z-NA (green) and ASFV structure protein P30 (red). Nuclei were stained by DAPI (blue). (F) Quantifying the mean fluorescence intensity (MFI) of Z-NA staining in ASFV-infected cells in C-E. Scale bar, 5 μm. (G) ASFV CN/GS/2018 (MOI = 1.0) infected PAMs were co-stained with ZBP1 and Z-NA antibodies. (H) Quantitative analysis of the co-localization between ZBP1 and Z-DNA was performed with ImageJ. (I) Immunoblot analysis of ZBP1, p-MLKL, and MLKL in nuclear and cytoplasmic fractions from ASFV-infected (MOI = 1.0) PAMs at the indicated times. Immunoblotting for β-actin and Lamin A/C was used to confirm the purity of cytoplasmic and nuclear fractions. (J) MA-104 cells were transfected with Myc-ZBP1^WT^ or ZBP1^ΔZα2^ and then infected with ASFV (MOI = 1.0) to analyze the ZBP1 localization at 16 hpi. ASFV infection was indicated by co-staining for p-MLKL in the same cells. Scale bars represent 5 μm (C, D, E, G, and J).

To investigate the relationship between ZBP1 and Z-DNA during ASFV infection, the subcellular localization of ZBP1 and Z-DNA in PAMs infected with ASFV was further analyzed by IFA. IFA results showed a distinct co-localization between ZBP1 and Z-DNA in the cytoplasm and nucleus of ASFV-infected PAMs (**Fig. 6G and 6H**). We also examined the subcellular location of ZBP1 following ASFV infection by separating the nucleus and cytoplasm components. The Western-blotting results show that ZBP1 preferentially upregulated in the nucleus at 8 hpi and subsequently appears in the cytoplasm at 12 hpi. And the MLKL also was phosphorylated initially in the nucleus and then in the cytoplasm of ASFV-infected PAMs (**Fig. 6I**), suggesting that the ZBP1-mediated necroptosis signaling was activated in both the cytoplasm and nucleus of ASFV-infected cells.

The Zα domain of ZBP1 is important for sensing accumulated Z-NA and its nuclear migration (29, 30). By reconstituting the Myc-tagged ZBP1 mutant plasmid lacking the Zα2 domain (Myc-ZBP1^ΔZα2^) that based on the Myc-tagged porcine wild-type ZBP1 plasmid (Myc-ZBP1^WT^) (**Fig. S3**), the role of the Zα2 domain on the ZBP1 nucleus localization after ASFV infection was further investigated. Given that overexpressed plasmids are difficult to transfect into primary macrophages, Myc-ZBP1^WT^ and Myc-ZBP1^ΔZα2^ plasmids were transfected into MA-104 cells (a passage cell line that can infect ASFV) and then infected with ASFV to observe their subcellular localization (31). The results showed that Myc-ZBP1^WT^ colocalized with p-MLKL in the nucleus and cytoplasm of ASFV-infected MA-104 cells. However, Myc-ZBP1^ΔZα2^ was detained in the cytoplasm of ASFV-infected MA-104 cells, similar to the subcellular localization of Myc-ZBP1^WT^ expressed in uninfected MA-104 cells. Consequently, the Zα2 domain of ZBP1 plays an important role in response to ASFV-induced nucleus localization of ZBP1 (**Fig. 6J**).

### ZBP1-mediated macrophage necroptosis facilitates the extracellular release of proinflammatory cytokines induced by ASFV infection

Necroptosis causes cell plasma membrane damage, further resulting in the extracellular release of cytoplasmic proinflammatory cytokines and DAMPs (17). The concentration of proinflammatory cytokines, including porcine TNF-α, IL-1β, IL-18, and IFN-γ in supernatants from ASFV-infected macrophages, were measured using ELISA kits. After infection with ASFV, the concentrations of porcine TNF-α and IL-1β were significantly elevated in supernatants of PAMs and BMDMs in a time-dependent manner (**Fig. 8A-8D**). However, the porcine IL-18 and IFN-γ levels do not change meaningfully in supernatants of cells infected with ASFV (**Fig. S4**).

**FIG 7.**
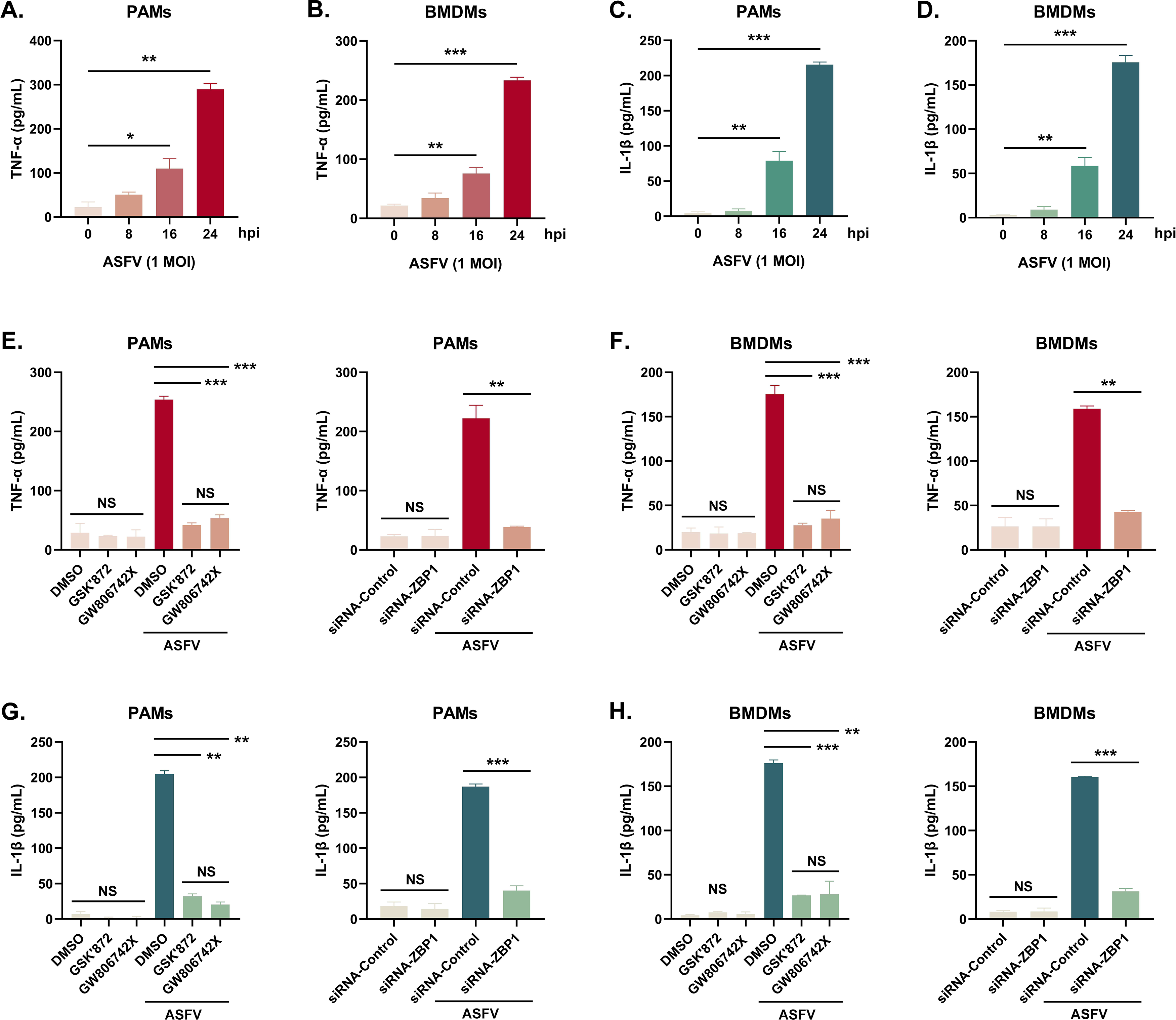
The extracellular release of TNF-α and IL-1β was restrained in ASFV-infected macrophages by inhibiting the ZBP1-mediated necroptosis. The TNF-α (A, C, E, F, I, and J) and mature IL-1β (B, D, G, H, K, and L) levels in the supernatants were determined by ELISA. PAMs (A, B, E-H) and BMDMs (C, D, I-L) were infected with ASFV at 1.0 MOI for indicated times. (E, G, I, and K) PAMs and BMDMs were treated with GSK’872 (1 μM) or GW806742X (1 nM) to inhibit necroptosis or were added to DMOS as a control. (G, H, K, and L) PAMs and BMDMs were transfected with siRNA-ZBP1 or siRNA-control before infecting with ASFV. Data shown are means ± SD. Statistical significance was analyzed by Student’s t-test. \**P* < 0.05; \*\**P* < 0.01; \*\*\**P* < 0.001; \*\*\*\**P* < 0.0001.

**FIG 8.**
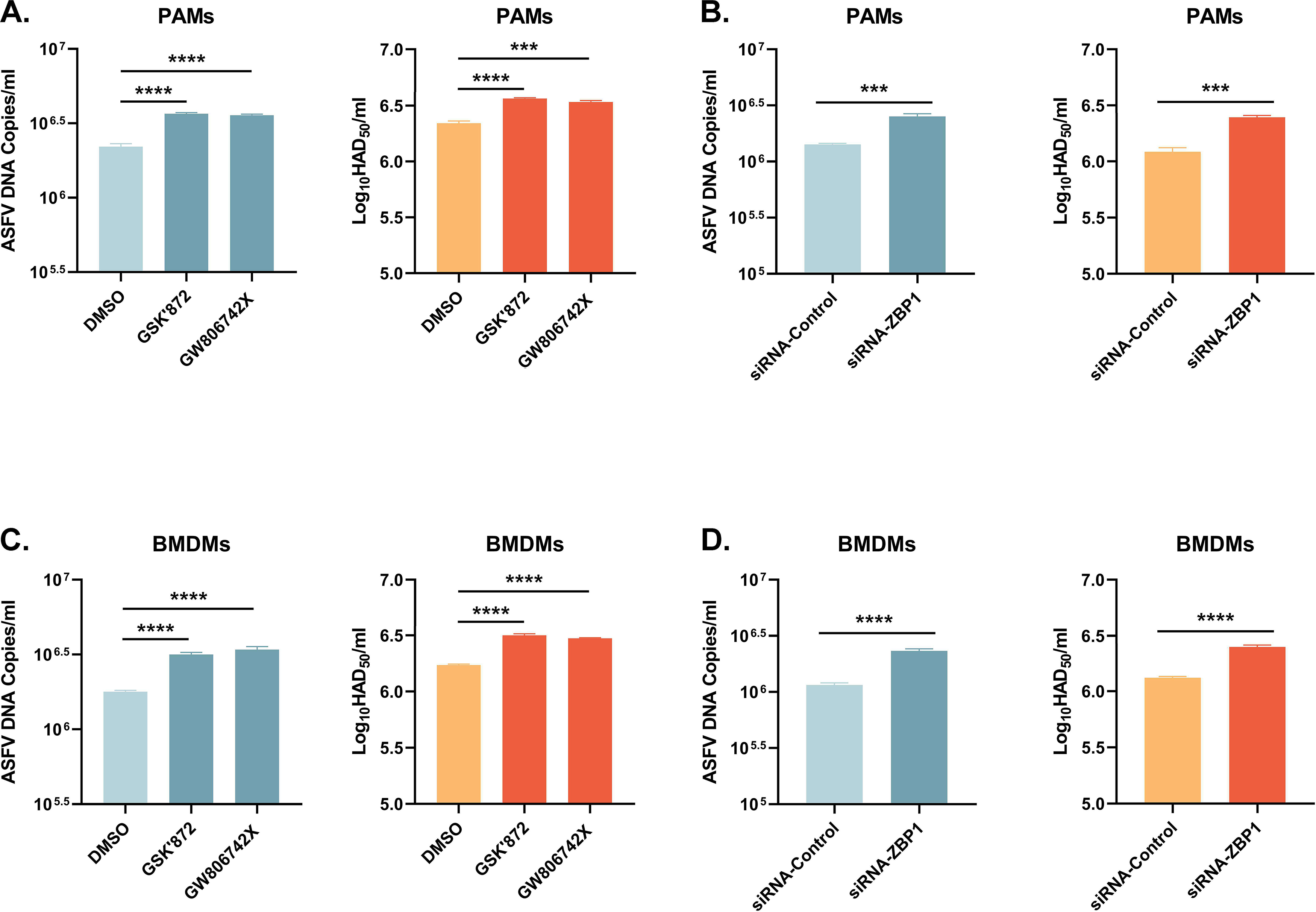
ASFV replication was boosted in PAMs and BMDMs, in which necroptosis was inhibited. PAMs (A and B) and BMDMs (C and D) were infected with ASFV at 1.0 MOI for 24 h. PAMs (A) and BMDMs (C) were treated with GSK’872 (1 μM) or GW806742X (1 nM) to inhibit necroptosis during infection. And PAMs (B) and BMMDs (D) were transfected with siRNA-ZBP1 before infecting with ASFV. ASFV genome copies and viral titer were determined by qRT-PCR and HAD_50_. Data shown are means ± SD. Statistical significance was analyzed by Student’s t-test. \**P* < 0.05; \*\**P* < 0.01; \*\*\**P* < 0.001; \*\*\*\**P* < 0.0001.

Furthermore, the effect of ZBP1-mediated necroptosis signaling on ASFV-induced release of TNF-α and IL-1β was investigated. After inhibiting the macrophage necroptosis using inhibitors or siRNA-ZBP1, TNF-α and IL-1β were severely dampened in supernatants of PAMs and BMDMs infected with ASFV (**Fig. 8E-8H**). Together, these results demonstrated that ZBP1-mediated necroptosis is the main cause of the extracellular release of proinflammatory cytokines in ASFV-infected macrophages.

### ZBP1-mediated necroptosis inhibits ASFV replication in macrophages

Necroptosis signaling induces severe inflammation during viral infection but has different effects on viral replication. Western blot results showed that the expression of ASFV P72 was upregulated in necroptosis-inhibited macrophages by treating inhibitors or transfecting siRNA-ZBP1. We finally examined the role of ZBP1-mediated macrophage necroptosis on ASFV replication. After treatment with inhibitors or siRNA, the ASFV genome copies and viral titer in PAMs and BMDMs infected with ASFV were determined by qRT-PCR and HAD_50_ assay. The results indicated that the ASFV genome copies and viral titer were markedly rise in necroptosis-inhibited PAMs and BMDMs (**Fig. 9**). Thus, our findings demonstrated that the ZBP1-mediated necroptosis signaling downregulatesASFV replication in macrophages.

**FIG 9.**
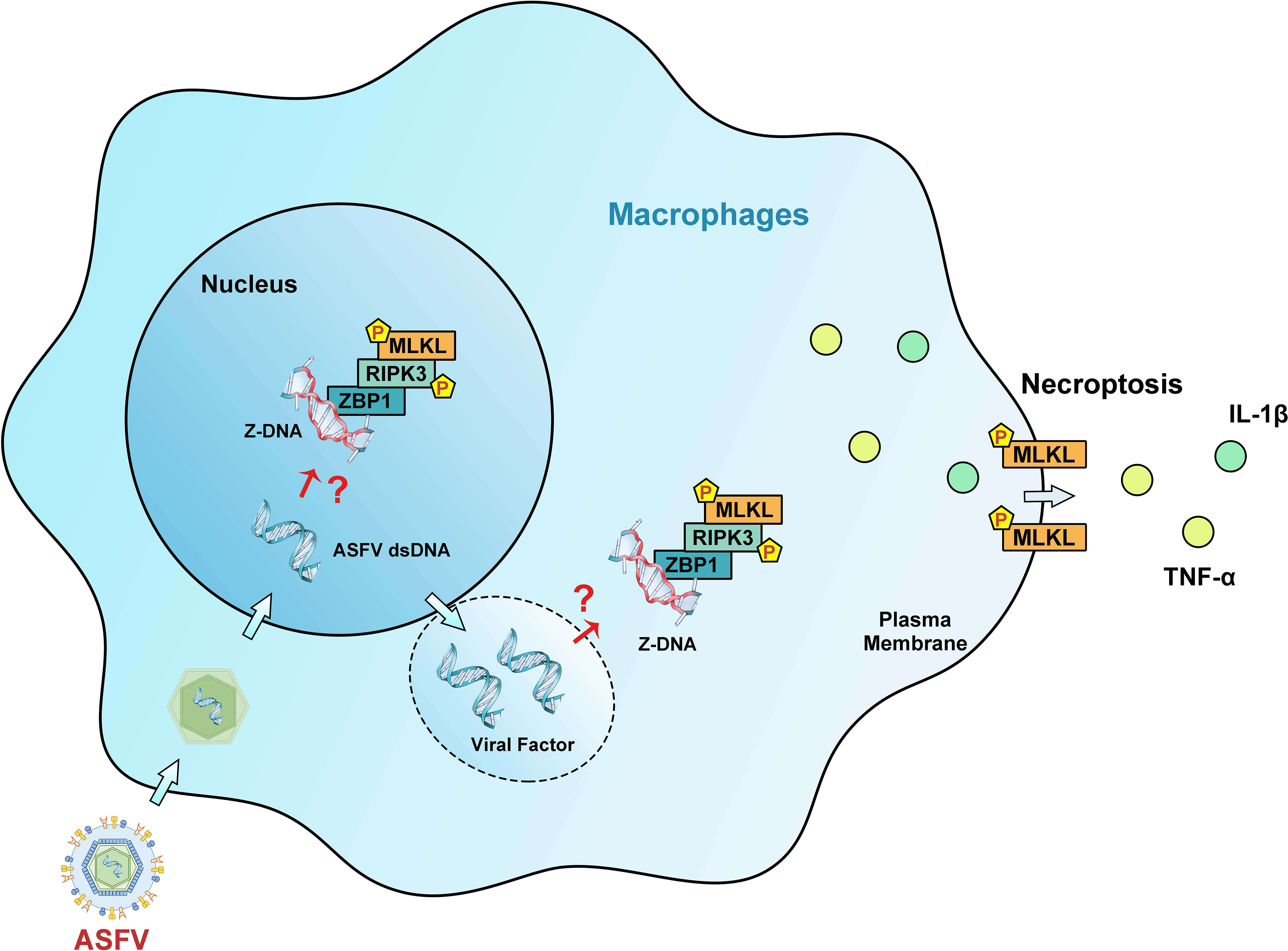
Schematic diagram of the mechanism of ZBP1-mediated necroptosis signaling in ASFV-infected macrophages. After invading macrophages, ASFV replicates initially in the nucleus and then in the cytoplasm of macrophages. Upregulated ZBP1 senses the accumulative Z-DNA in the nucleus and cytoplasm of ASFV-infected macrophages and recruits RIPK3 and MLKL to consist of the ZBP1 necrosome. The ZBP1-mediated necroptosis signaling promotes the MLKL-pores formation on the cytoplasmic membrane, resulting in the extracellular release of cytoplasmic proinflammatory cytokines. Finally, ZBP1-mediated necroptosis restricts the ASFV replication in macrophages.

## Discussion

Macrophages, highly plastic cells in the host immune system, are important for activating innate and adaptive immunity against microbes and causing various pathologies during infection (32). The interactions between ASFV and porcine macrophages play a critical role in ASFV-induced pathological lesions, whereas little is known about the interaction and pathogenic mechanism (33). We showed that ASFV infection activates the necroptosis signaling in the porcine spleen, lung, liver, and kidney (**Fig. 1**). And porcine macrophage necroptosis was induced by ASFV infection in *vitro* (**Fig. 2**). Hemorrhagic or hyperaemic splenomegaly is the most obvious lesion in ASFV-infected pigs (34). The histopathological of the ASFV-infected spleen mainly renders as a hyperaemic red pulp, even filled with red blood cells (33). The macrophage population of porcine splenic red pulp has predominantly anchored a mesh of fibers in the splenic cords, surrounded by smooth muscle cells (35). The macrophage death in the red pulp will cause damage to intercellular junctions between the smooth muscle cells and further bare the basal lamina, followed by the activation of the coagulation cascade, platelet aggregation, and fibrin deposition, finally accumulating red blood cells within the splenic cords (33, 36). Thus, macrophage necroptosis may explain why ASFV infection results in hemorrhagic or hyperaemic splenomegaly.

Mechanically, TNF-α expression was induced by ASFV (7). Overexpressing ASFV A179L enhanced the TNF-α-induced and RIPK1-dependent necroptosis (23), TLR3 and TLR4 were activated to induce the IL-1β expression in ASFV-infected PAMs (24). However, our results confirmed that ASFV-induced macrophage necroptosis is independent of TNF-α-RIPK1 and TLR-TRIF pathways (**Fig. 3 and 4**). RIPK1 is an important adapter protein for signaling transduction, including necroptosis, apoptosis, pyroptosis, and MAPK signaling (37). Many viral proteins target RIPK1 to suppress the RIPK1-mediated signaling. Murine cytomegalovirus (MCMV) encoded M45 and herpes simplex virus 1 (HSV-1) encoded ICP6 both inhibit the necroptosis signaling by disrupting the RIPK1-RIPK3 interaction (38-40). And epstein-barr virus (EBV) encoded LMP1 interacts and ubiquitinated RIPK1 to suppress necroptosis (41). Thus, proteins encoded by ASFV may target RIP1 to inhibit TNF-α-induced necroptosis. In addition, ASFV I329L has been reported as an inhibitor of TLR3 signaling by targeting TRIF (42). That may also be responsible for the suppression of TLR-TRIF pathway-dependent necroptosis.

ZBP1 was originally identified as an innate immune sensor that activates interferon (IFN) responses and NF-κB signaling (43). Recently, ZBP1 has been reported as the master protein inducing PANoptosis (pyroptosis, apoptosis, and necroptosis) (29). Our past and present results confirmed that ZBP1 expression was upregulated in host macrophages upon ASFV infection (**Fig. 5A-C**) (25, 26). The upregulated ZBP1 interacts with RIPK3 and MLKL to consist of the ZBP1-RIPK3-MLKL necrosome and activate macrophage necroptosis during ASFV infection (**Fig. 5D-H**).

Virus infection-induced ZBP1 activation needs ZBP1 to sense the virus-derived Z-RNA (30, 44, 45). Differently, Z-DNA sequences were predicted to be present in the ASFV genome using the Z-HUNT3 program (**Fig. 6B**). Further experiments confirmed that Z-DNA signals, but not Z-RNA signals, did accumulate in both the nucleus and cytoplasm of ASFV-infected cells (**Fig. 6C-F**). Past studies have confirmed that early replication of ASFV occurs in the nucleus and subsequently replicates in the perinuclear viral factory (46, 47), further enhancing the credibility of this hypothesis that viral dsDNA generated from ASFV replication forms Z-DNA, although this needs to be confirmed by more experiments.

The subcellular distribution of ZBP1-mediated necroptosis signaling mainly depends on the ZBP1 recognization for its Z-NA ligands. Influenza A virus (IAV) replication generates Z-RNA in the nucleus so that ZBP1 initiates necroptosis activation in the nucleus and results in nuclear envelope disruption leakage of DNA into the cytosol (30). However, ZBP1 recognized viral Z-RNA and activated necroptosis in the vaccinia virus (VACV) cytoplasm or SARS-CoV-2 infected cells (44, 45). Past studies have confirmed that early replication of ASFV occurs in the nucleus and subsequently replicates in the perinuclear viral factory (46, 47). Our finding demonstrated the co-localization between ZBP1 and Z-DNA in the nucleus and the cytoplasm of ASFV-infected cells (**Fig. 6G-H**), activated the nucleus and cytoplasmic necroptosis signaling (**Fig. 6I**). However, in ASFV-infected cells, we did not observe the clear leakage of nuclear chromatin and DNA into the cytosol, similar to the IAV-infected cells. The effect of nucleus necroptosis signaling in ASFV-infected cells is currently unknown.

The virus-infected monocyte-macrophages will extensively express and secret proinflammatory cytokines, including TNF-α, and IL, described as cytokines storm (8, 22). Our findings indicated that ASFV infection induces the massive secretion of TNF-α and IL-1β in macrophages, but IFN-γ and IL-18 do not change (**Fig. 7**). Either pyroptosis or necroptosis contributes to the extracellular release of cytoplasmic proinflammatory cytokines by damaging the cytoplasm membrane. We found that the extracellular levels of TNF-α and IL-1β were almost completely blocked and dampened when ASFV-induced macrophage necroptosis was inhibited (**Fig. 7**). During IAV and SARS-CoV-2 infection, infected cells released proinflammatory cytokines that were significantly alleviated only when pyroptosis and necroptosis were both inhibited. This difference may be attributed to the GSDMD inactivation caused by ASFV pS273R, causing ASFV-induced proinflammatory cytokines only to be released from macrophages via MLKL-pores (15).

ZBP1-mediated necroptosis effectively restricts the ASFV infection in macrophages (**Fig. 8**). However, the ZBP1-dependent PANoptosis-mediated cytokines storm usually results in severe organ dysfunction and clinical symptoms. The inflammatory cell infiltration and alveolar septa expansion caused by SARS-CoV-2 infection were meaningfully alleviated after knocking down RIPK3 (45, 48). Necroptotic cells act as impactful immunogenic to recruit and activate neutrophils, and bronchioalveolar necroptosis is also the reason for IAV-induced acute respiratory distress syndrome (ARDS) (49, 50). Hemorrhagic lesions caused by ASFV are primarily due to the release of cytokines from infected macrophages rather than direct damage to endothelial cells (36). We also confirmed that ASFV activated necroptosis signaling in porcine lungs and PAMs, which may be related to respiratory distress and severe hemorrhages in the septa and the alveolar spaces for ASFV-infected pigs (9, 33). Massive destruction of the lymphoid organs and tissues, including the spleen and lymph nodes, are the most important features in ASFV pathology. Many B and T lymphocytes undergo cell death in acute ASFV infection (51, 52). In immune organs, signaling interaction between lymphocytes and macrophages is key to triggering the adaptive immune response. However, during ASFV infection, ZBP1-mediated necroptotic macrophages release TNF-α and IL-1β that will cause lymphocyte apoptosis (53). ASFV-induced macrophage necroptosis may be a key pathway by which ASFV disrupts the porcine adaptive immune system.

In conclusion, this work demonstrates that ASFV can induce macrophage necroptosis by promoting ZBP1 necrosome activation. It also uncovers the mechanism by which Z-DNA in ASFV-infected cells is recognized by ZBP1 to induce necrosome assembly. And ZBP1-mediated necroptosis signaling causes the release of proinflammatory cytokines and inhibits the ASFV replication in macrophages. Regrettably, whether ZBP1-mediated macrophage necroptosis is beneficial or harmful for host immunity against ASFV infection still requires further animal experiments to elucidate. Together, these findings extend our knowledge about the interaction between ASFV and macrophage necroptosis and shed light on the pathogenesis of ASFV.

## Materials and methods

### Animal experiments and ethics statement

Animal experiments were conducted at the Lanzhou Veterinary Research Institute (LVRI) of the Chinese Academy of Agricultural Sciences (CAAS, China). Four landrace pigs aged two-month-old and weighing approximately 25 Kg were obtained from a licensed livestock farm, with each pig being antigenically or serologically negative for ASFV, porcine reproductive and respiratory syndrome virus, pseudorabies virus, or porcine epidemic diarrhea virus, and classic swine fever virus. In this experiment, two random pigs were injected with ASFV/CN/GS/2018 (1.0 HAD50) as the ASFV infection group. And other two pigs were injected with phosphate-buffered saline as a mock infection group. All pigs were euthanized when critically ill or on 10 the day to collect the required tissue samples.

All animal experiments (including the euthanasia procedure) were performed in strict accordance with the Animal Ethics Procedures and Guidelines of the People’s Republic of China and Accreditation of Laboratory Animal Care International and the Institutional Animal Care and Use Committee guidelines (License No. SYXK [GAN] 2014–003). All experiments involving live ASFV have performed in the biosafety level 3 facilities of Lanzhou Veterinary Research Institute, Chinese Academy of Agricultural Sciences.

### Cell culture and virus

Porcine alveolar macrophages (PAMs) were prepared using bronchoalveolar lavage as described previously and cultured in Roswell Park Memorial Institute (RPIM) 1640 medium (Gibco, USA) containing 10% porcine serum (Gibco, USA). Porcine bone marrow cells were harvested from the hip bone and femur of 8-week-old pigs. Harvested cells were cultured in RPIM 1640 medium containing 20% porcine serum and 10 ng/mL recombinant porcine granulocyte-macrophage colony-stimulating factor (GM-CSF, R&D Systems, UK) to differentiate mature bone marrow-derived macrophage (BMDMs) after 7 days. BMDMs were cultured in RPIM 1640 medium containing 20% porcine serum. Microbiological associates-104 (MA-104) cells were purchased from China Center for Type Culture Collection (Wuhan, China) and were cultured in Dulbecco’s modified Eagle medium (Gibco, USA) with 10% fetal bovine serum (FBS, VivaCell, New Zealand). All cells were cultured in a humidified atmosphere containing 5% CO_2_ at 37℃.

The CN/GS/2018 ASFV, a genotype II strain, was isolated and preserved by the Lanzhou Veterinary Research Institute, Chinese Academy of Agricultural Sciences. Viral titration of the strain was mensurated using the hemadsorption (HAD) assay and presented in HAD_50_ per milliliter.

### Antibodies and reagents

Anti-B646L (ASFV P72 structure protein) mouse polyclonal antibody was prepared and provided by LVRI, CAAS. Anti-CP204L (ASFV P30 structure protein) mouse monoclonal antibody was presented by the College of Veterinary Medicine, Nanjing Agricultural University (Nanjing, China). Rabbit anti-MATR3 (ab151714) antibody was purchased from Abcam. Mouse anti-ZBP1 monoclonal antibody (sc-271483) was purchased from Santa Cruz Biotechnology, Inc (Texas, USA). Rabbit anti-Z-NA antibody (Z22) was purchased from Absolute Biotech Company (Oxford, UK). The following rabbit monoclonal antibodies were used: anti-Lamin A/C (4777S), anti-RIPK3 (13526S), anti-p-RIPK3 (91702S), anti-p-MLKL (37333S), anti-Myc-Tag (2272S), Alexa Fluor 488 anti-rabbit IgG (4416S), and Alexa Fluor 594 anti-mouse IgG (8890S) were purchased from Cell Signaling Technology (Massachusetts, USA); Anti-MLKL rabbit monoclonal antibody (ET1601-25) was purchased from HuaAn McAb Biotechnology Co, Ltd (Hangzhou, China). Anti-β-actin mouse monoclonal antibody (66009-1-Ig), Horseradish peroxidase (HRP)-conjugated goat anti-mouse IgG (H+L; SA00001-1), HRP -conjugated goat anti-rabbit IgG (H+L; SA00001-2) were purchased from Proteintech Group Co., Ltd. (Chicago, IL, USA). Protein G Sepharose (17061801) was purchased from GE Healthcare Bio-sciences AB (Pittsburgh, PA, USA). HRP-conjugated goat anti-mouse IgG (SE131) was purchased from Beijing Solarbio Science And Technology Co., Ltd (Beijing, China).

Recombinant human IFN-β (300-02BC) was purchased from Peprotech (US). LPS (L8880) was purchased from Solarbio Life Science (Beijing, China). Recombinant porcine GM-CSF protein (711-PG-010) was purchased from R&D System (Minnesota, USA). SM-164 (HY-15989), Z-VAD-FMK (HY-16658B), Necrostatin-1 (HY-15760), GSK’872 (HY-101872), GW806742X (HY-112292), and Pepinh-TRIF (HY-P2565) were purchased from MedChemExpress (New Jersey, USA). Polyplus jetPRIME (PT-114-15) transfection reagent was purchased from Polyplus-Transfection SA (Strasbourg, France). Lipofectamin RNAiMAX transfection reagent was purchased from Thermo Fisher Scientific (California, USA).

### Tissue protein extraction

100 mg tissues were added to 800 μl of strong RIPA lysis buffer supplemented with protease and phosphatase inhibitor in 1.5 ml tubes. Add sterile steel balls to each tube and fully grind the tissue using a TIANGEN H24 grinding homogenizer (Beijing, China). Tissues protein were separated by centrifuging the tissue lysates (12000 rpm, 10 min). Protein concentration was detected by BCA kit (Thermo Fisher Scientific, 23227), then was quantified to 5 μg/μl and further Western blot assay.

### Plasmids construction and transfection

Plasmids encoding porcine ZBP1 (Gene ID: 100144524) and ZBP1^ΔZα2^ (deleting its Zα2 domain) were constructed by cloning the synthesized sequence into pcDNA3.1 with Myc tag fused to the 3′ end. MA-104 cells were cultured in 12-well plates and transfected with 3 μg plasmids per well using Polyplus jetPRIME transfection reagent. After 8 h, cells were replaced with fresh DMEM culture medium. At 24 h post-transfection, cells were further treated or infected with ASFV.

### RNA interference

The siRNA duplexes targeting porcine ZBP1 were chemically synthesized by Gene-Pharma. ZBP1-specific siRNA duplexes were transfected into PAMs or BMDMs with RNAiMAX transfection reagents according to the manufacturer’s instructions. After 5 h, the fresh medium replaced the medium containing transfection reagents, and cells were further infected for ASFV. The sequences of ZBP1-specific siRNA oligonucleotide are as follows: 5′-CCAGAGGCUUCCAUUGAUATT-3′.

### Real-time qPCR for determination of ZBP1 gene expression

Total RNA was extracted from PAMs using the TRIzol reagent and was reverse transcribed using the PrimeScript RT kit (TaKaRa). qPCR was performed using the PowerUp SYBR Green Master Mix on the ABI StepOnePlus system. All data were analyzed using the StepOnePlus software, and the relative mRNA level of ZBP1 was normalized based on the porcine glyceraldehyde 3-phosphate dehydrogenase (GAPDH) mRNA level. Furthermore, the relative mRNA level of ZBP1 was determined based on the comparative cycle threshold (2^−ΔΔCT^) method. The primer sequences are as follows:

Porcine ZBP1-F: 5′-GAAGGGACTGAACAACTGCA-3′

Porcine ZBP1-R: 5′-CGTCGGAGGGAGGTAGAA-3′

Porcine GAPDH-F: 5′- ACATGGCCTCCAAGGAGTAAGA −3′

Porcine GAPDH-F: 5′- GATCGAGTTGGGGCTGTGACT −3′

### Real-time qPCR detecting ASFV genome copies

ASFV genomic DNA in infected cells was extracted using QIAamp DNA Mini Kits (Qiagen, Germany). RT-qPCR was performed using the Pro Taq HS premix Probe qPCR kit (Accurate Biology, China) and the QuantStudio5 system (Applied Biosystems, USA). The targeting gene for amplification of the ASFV genome was the conserved B646L gene segment, using the following primers:

ASFV-P72-F: 5′-GATACCACAAGATCAGCCGT-3′

ASFV-P72-R: 5′-CTGCTCATGGTATCAATCTTATCGA-3′

TaqMan probe: 5′-CCACGGGAGGAATACCAACCCAGTG-3′

The amplification conditions used to detect ASFV genome copies were as follows: preheating at 95°C for 30 s; 40 cycles at 95°C for 5 s; and annealing at 58°C for 30 s; elongation (72°C). The copy numbers of the ASFV genome were calculated using an established standard curve and presented in genome copies per milliliter.

### Viral titration

The ASFV CN/GS/2018 strain was quantified using hemadsorption (HAD) assays with minor modifications. Briefly, PAMs and porcine red blood cells were seeded in 96-well plates. The samples containing ASFV were then added to the plates and titrated in 10-fold serial dilutions. HAD was observed at day 7 post-inoculation, and 50% HAD doses (HAD_50_) were calculated according to the Reed–Muench method.

### Co-immunoprecipitation (Co-IP)

ASFV-infected PAMs were lysed in NP-40 lysis buffer supplemented with 1 mM phenylmethanesulfonyl fluoride. Cell lysates were incubated on ice for 30 min and briefly sonicated, then high-speed centrifugation (12000 rpm, 10 min) at 4℃. After saving 10% of the total cell lysate for input, the extracts were divided into two parts for immunoprecipitation with anti-ZBP1 or mouse IgG at 4°C overnight. Subsequently, protein G agarose beads (Roche) were added to the cell lysate and incubated for 3 h at 4°C. Finally, beads were centrifuged (8000 rpm, 30 s) and washed thrice with PBS buffer. The protein-antibody-beads complex was added to the SDS-loading buffer and further detected by immunoblotting.

### Immunoblotting analysis

The proteins with different molecular weights were separated by 8% using SDS-PAGE (80 V, 30 min; 120 V, 60 min) and then transferred to a nitrocellulose membrane (100 V, 90 min). The protein-loaded nitrocellulose membrane was blocked with bovine serum albumin (5%) for 1 h at normal temperature and incubated with the designated antibodies at 4°C overnight. HRP-conjugated anti-mouse/rabbit IgG was incubated to bind the designated antibodies. After developing an electrochemiluminescence solution, the immunoblotting images were captured by the Odyssey infrared imaging system.

### Indirect immunofluorescence assay

Cells were plated on 20 mm glass-bottom cell culture dishes in diameter (NEST) and allowed to adhere for at least 8 h before the next experiments. Following ASFV infection or other treatments, cells were fixed in freshly-prepared 4% (w/v) paraformaldehyde, permeabilized in 0.2% (v/v) Triton X-100, blocked with 5% BSA (Sigma), and incubated overnight with primary antibodies at 4℃. Then, dishes were incubated with fluorophore-conjugated secondary antibodies for 2 h at room temperature. Following three additional washes in PBS, the cell nucleus was stained with 4-methyl-6-phenylindole for 10 min and imaged by confocal microscopy on a Leica SP8 instrument. Fluorescence intensity was quantified using Leica LAS X software.

### Cytotoxicity assay

PAMs and BMDMs were cultured in 48-well Cell culture plates. Discard the supernatant after cell adhesion, and cells were infected with ASFV. After 2 h, the viral fluid was replaced with a fresh 1640 culture medium containing 2% FBS. At the indicated times, the cell supernatant was collected and detected immediately. Cytoplasmic membrane damage was detected by measuring cytoplasmic lactate dehydrogenase (LDH) release using Promega CytoTox 96^®^Non-Radioactive Cytotoxicity Assay kit (J2380) according to the manufacturer’s instructions. Luminescence was measured on a Synergy HT Multi-Detection microplate reader (Bio-Tek). Relative LDH release was calculated as the ratio of LDH release of experimental samples to maximal release.

### ELISA Assay

PAMs and BMDMs were cultured in 48-well Cell culture plates. Discard the supernatant after cell adhesion, and cells were infected with ASFV. After 2 h, the viral fluid was replaced with a fresh 1640 culture medium containing 2% FBS. At the indicated times, the cell supernatant was collected and detected immediately. The concentrations of porcine TNF-α (RAB047), IFN-γ (RAB0226), IL-1β (ESIL1B), and IL-18 (BMS672TEN) in supernatants were measured using the ELISA Kit (Sigma-Aldrich) according to the manufacturer’s instructions. The concentrations of porcine IL-1β (ESIL1B) and IL-18 (BMS672TEN) in culture supernatants were measured using the ELISA Kit (InvitrogenTM) according to the manufacturer’s instructions.

### Z-DNA sequences prediction

The Z-forming potential of the genome sequence of ASFV CN/GS/2018 was determined by the ZHUNT3 program (https://github.com/Ho-Lab-Colostate/zhunt) using the following command line "zhunt3nt 12 6 24". Data are presented using Microsoft Excel.

### Statistical analysis

All in vitro experiments were performed at least thrice. Data are presented as the means ± standard deviations (SDs). The statistical significance between groups was determined using the *t*-test with GraphPad Prism v.8 (San Diego, CA, USA). \**P* < 0.05, \*\**P* < 0.01, \*\*\**P* < 0.001, \*\*\*\**P* < 0.0001 was considered statistically significant. NS: no significant difference.

## Acknowledgments

This work was supported by grants from the National Key R&D Program of China (2021YFD1801300); This work was supported by the major science and technology project of Gansu Province, China (22ZD6NA001, 22ZD6NA012); The Research funding from National Swine Technology Innovation Center of China (NCTIP-XD/C03, CARS-35). This work was supported by the Institute of Animal Health, Guangdong Academy of Agricultural Sciences of China (2022SDZG02). The authors would like to thank all the editors and reviewers for their valuable comments and suggestions that helped improve the quality of this manuscript.

---

**FIG S1. The proliferation kinetics of ASFV in different cells.** PAMs (A), BMDMs (B), and MA-104 cells (C) were infected with ASFV (MOI = 0.01), and the viral titers at 12, 24, 48, 72, and 96 hpi were determined using the HAD_50_ method. Data are presented as mean ± SD of three independent experiments.

**FIG S2. The upregulated ZBP1 in the spleen, lung, liver and kidney derived from pigs infected with ASFV.**

**FIG S3. The construction and expression of Myc-tagged porcine ZBP1^WT^ and ZBP1^ΔZα2^ plasmids.**

**FIG S4. The IFN-γ and IL-18 concentration in the supernatants of PAMs or BMDMs after ASFV infection.**

**Supplementary Table 1.** Z-HUNT3 analysis results on ASFV CN/GS/2018 genome.

